# Description of *Ruminococcus catenae* SW178 sp. nov., a new anaerobic species isolated from feral chicken

**DOI:** 10.1101/777680

**Authors:** Supapit Wongkuna, Sudeep Ghimire, Surang Chankhamhaengdecha, Tavan Janvilisri, Joy Scaria

## Abstract

A Gram-positive, obligately anaerobic coccobacillus, with the white raised circular colony was isolated from the cecum of feral chickens in Brookings, South Dakota, USA. The 16S rRNA gene sequence analysis suggested that the closest species to strain SW178 was *Ruminococcus torques* ATCC 27756^T^ (96.94% similarity) that belongs to the family *Lachnospiraceae*. The genome of strain SW178 is 3.18 Mbp with G+C content of 46.9 mol%. Based on the phylogenetic and phenotypic comparison, we propose that strain SW178 be assigned to the genus *Ruminococcus* as a novel species, for which the name *Ruminococcus catenae* is proposed. The type strain is SW178 (= CCOS 1886 ^T^, =DSM 109242^T^).

## Introduction

The bacterial community in the gastrointestinal tract of animals provide gut health benefits such as development of immune system, pathogen exclusion and digestion of polysaccharides (1, 2). Recent advancements in sequencing technology have helped to define overall gut microbiome composition of humans animals using metagenomics (3, 4). However, isolation and cultivation of gut microbiota composition are crucial to determine the phenotypic characteristics of individual species and to identify ecological interactions between microbes and the host (5, 6). Availability of cultured diversity also helpful for the development of probiotics. Herein, we report the isolation and taxonomic characterization of a new anaerobic bacterium from the cecum of feral chicken. We propose *Ruminococcus catenae* sp. nov. strain SW178 (= CCOS 1886 ^T^, =DSM 109242^T^) as a new member of genus *Ruminococcus.*

## Isolation and Ecology

Strain SW178 was isolated from the cecum of feral chicken. For cultivation, fresh cecal content was transferred within 10 minutes of the collection into an anaerobic workstation (Coy Labs, USA) containing 85% nitrogen, 10% hydrogen and 5□% carbon dioxide. Modified Brain Heart Infusion (BHI-M) medium containing 37 g/L of BHI, 5 g/L of yeast extract, 1 ml of 1 mg/ml menadione, 0.3 g of L-cysteine, 1 ml of 0.25 mg/L of resazurin, 1 ml of 0.5 mg/ml hemin, 10 ml of vitamin and mineral mixture, 1.7 ml of 30 mM acetic acid, 2 ml of 8 mM propionic acid, 2 ml of 4 mM butyric acid, 100 µl of 1 mM isovaleric acid, and 1% of pectin and inulin was used for isolation. The strain was maintained in BHI-M medium and stored with 10% (v/v) Dimethyl Sulfoxide (DMSO) at -80° C. Reference strain *R. torques* ATTCC 27756^T^, obtained from the American Type Culture Collection (ATCC), was maintained under the same conditions.

## 16S rRNA phylogeny

Genomic DNA of strain SW178 was extracted using a DNeasy Blood & Tissue kit (Qiagen), according to the manufacturer’s instructions. The 16S rRNA gene sequence was amplified using universal primer set 27F (5’-AGAGTTTGATCMTGGCTCAG-3’; Lane et al., 1991) and 1492R (5’-ACCTTGTTACGACTT-3’; Stackebrandt et al., 1993) (7, 8), and sequenced using a Sanger sequencer (ABI 3730XL; Applied Biosystems). The 16S rRNA gene sequence of SW165 was then compared with closely related strains from the GenBank (www.ncbi.nlm.nih.gov/genbank/) and EZtaxon databases (www.ezbiocloud.net/eztaxon) (9). Alignment and phylogenetic analysis were conducted using MEGA7 software (10). Multiple sequence alignments were generated using the CLUSTAL-W (11). The phylogeny was constructed using the maximum-likelihood (ML) (12), maximum-parsimony (MP) (13), and neighbour-joining (NJ) (14) methods. The distance matrices were generated according to Kimura’s two-parameter model. Bootstrap resampling analysis of 1000 replicates was performed to estimate the confidence of the tree topologies. Based on 16S rRNA gene sequencing, the closest taxon related to SW178 was *Ruminococcus torques* ATCC 27756^T^ with 96.94% similarity. The results of phylogenetic tree analysis showed that SW178 clustered within the *Merditerraneibacter* (*Ruminococcus*) indicating that SW178 represents a novel member of this genus (Fig. 1).

**Fig. 1.**
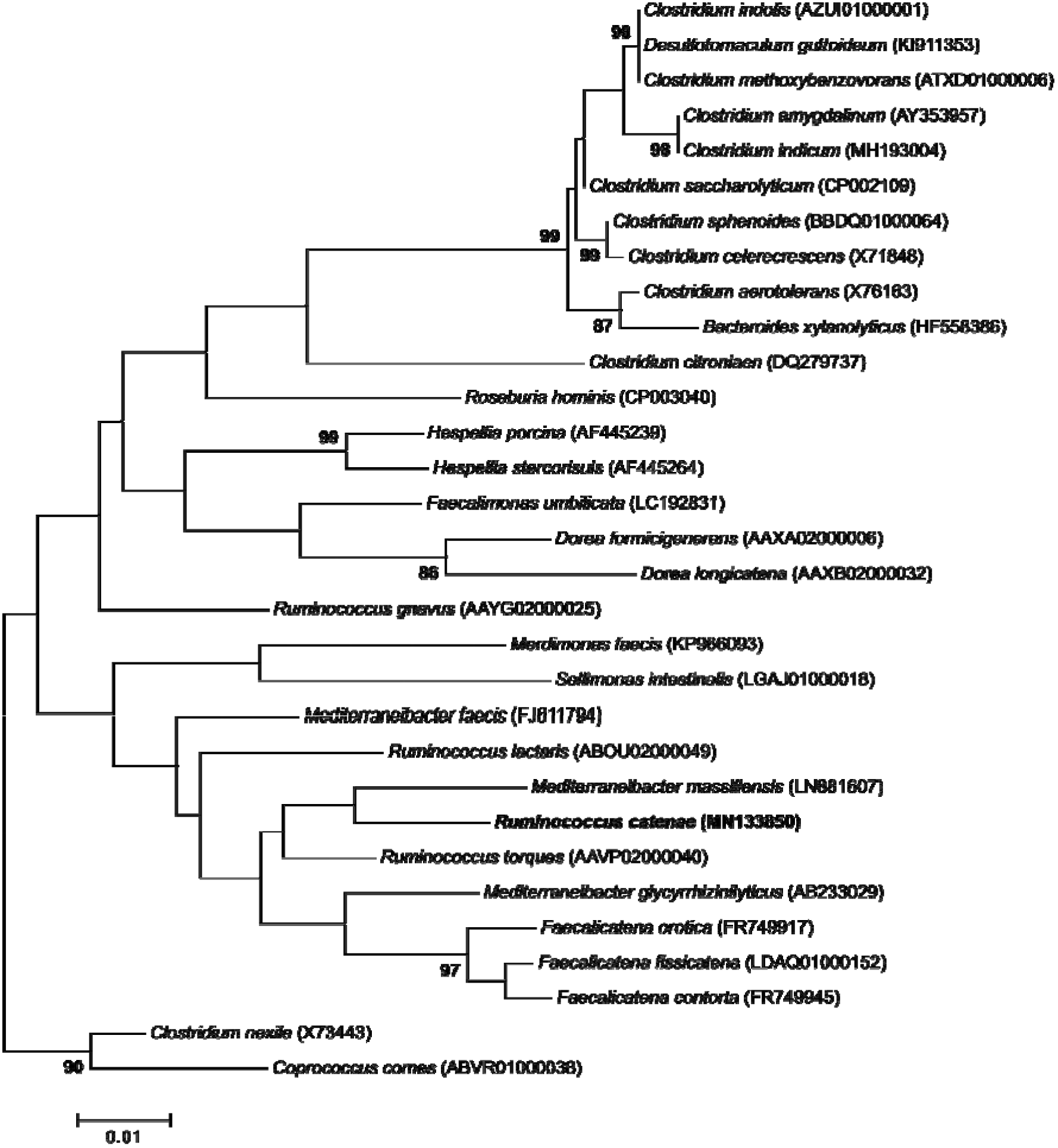
Maximum likelihood based 16S rRNA gene phylogeny, showing the phylogenetic position of *R. catenae* sp. nov. (= CCOS 1886 ^T^, =DSM 109242^T^) with closely related taxa in the family *Lachnospiraceae*. GenBank accession numbers of the 16S rRNA gene sequences are given in parentheses. Bootstrap values (based on 1000 replications) greater than or equal to 70 % are shown in percentages at each node. Bar, 0.01 substitutions per nucleotide position.

**Fig. 2.**
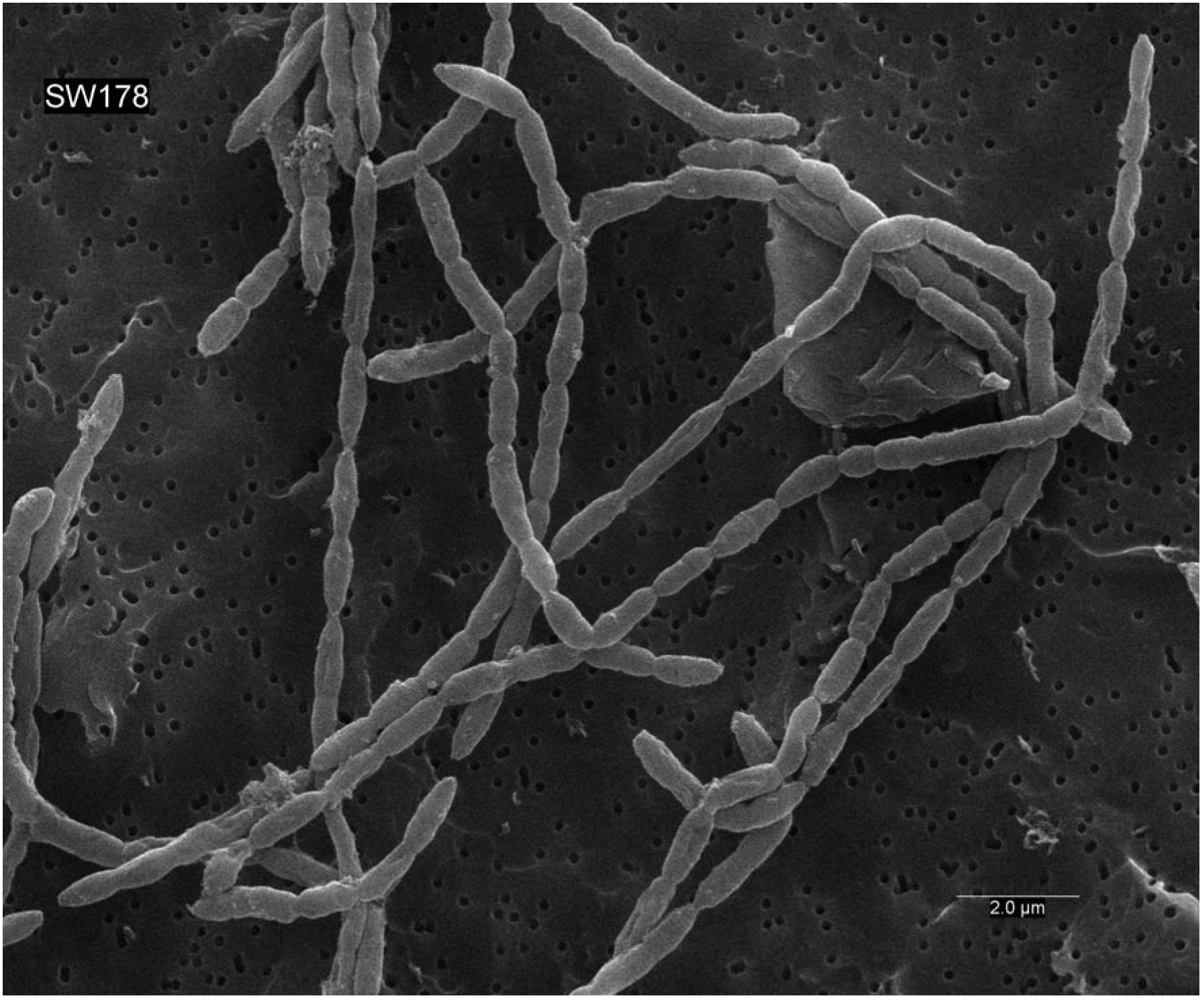
Scanning electron micrograph of cells of *Ruminococcus catenae* sp. nov. Cells were anaerobically cultured for 1 day at 37° C in BHI-M medium. Bar, 10.0 μm (a) and 2.0 μm (b).

## Genome features

The whole genome sequencing of strain SW178 was performed on Illumina MiSeq sequencer with MiSeq Reagent Kit V3 chemistry. The reads were assembled using Unicycler that builds an initial assembly graph from short reads using the de novo assembler SPAdes 3.11.1 (15). The quality assessment for the assemblies was performed using QUAST (16).The draft genome sequence of strain SW451 has a total length of 3.18 Mbp. The genomic G+C content of SW178 was 46.9□mol%. In addition, the average nucleotide identity (ANI) was calculated between SW178 and the closely related taxa using OrthoANI software (17). The ANI values were significantly less than the proposed ANI cutoff of 95–96□% (18) demonstrating strain SW178 as a novel species within the genus *Ruminococcus*.

## Physiology and Chemotaxonomy

Colony morphology was determined after 2-3 days of incubation on BHI-M agar plates. Gram-staining was performed using a Gram-Straining kit (Difco), according to the manufacturer’s instructions. Cell morphologies of cultures during exponential growth were examined by scanning electron microscopy (SEM). Aerotolerance was examined by incubating cultures for 2 days under aerobic and anaerobic conditions. Growth of strain SW178 at 4, 20, 30, 37, 40 and 55° C was observed. To determine pH range, the pH of the medium was adjusted to pH□4–9 with sterile anaerobic solutions of 0.1□M HCl and 0.1M NaOH. Motility of this microorganism was determined using motility medium with triphenyltetrazolium chloride (TTC) (19). The growth was indicated by the presence of red color, reduced form of TTC after it is absorbed into the bacterial cell wall. Other biochemical tests; utilization of various substrates and enzyme activities were determined using the AN MicroPlate (Biolog) and API ZYM (bioMérieux) according to the manufacturer’s instructions. For cellular fatty acid analysis, strain SW451 was cultured in BHI-M medium at 37□°C for 24□h under anaerobic condition. Cellular fatty acids were obtained from cell biomass and analyzed by GC (Agilent 7890A) according to the manufacturer’s instruction of Microbial Identification System (MIDI) (20).

Cells of the strain SW178 were Gram-positive coccobacilli with 0.5–1.0µm (Fig. 3). Colonies on BHI-M agar were white, raised and entire edge with 0.2-0.5 cm in diameter. Strain SW178 grew between 37° C and 45° C with optimum growth at 45°C. The optimum pH for the growth was 7.5, and growth was observed at pH□6.5–7.5. The strain could grow only under anaerobic condition indicating an obligately anaerobic bacterium. Based on the results obtained in Biolog tests, strain SW178 utilized dextrin, L-fucose, D-galacturonic, α-D-Glucose, L-rhamnose, and D-sorbitol. The strain SW178 exhibited positive reactions for esterase (C4), esterase lipase (C8), β-galactosidase, α-glucosidase and β-glucosidase on API ZYM plate (Table 1). The predominant cellular fatty acids of the strain SW178 included C_14□:□0_ (28.71%), C_16□:□0_ (19.55%) and C_16□:□0_ DMA (12.55%), in which differs from the reference strain (Table 2).

**Table 1.**
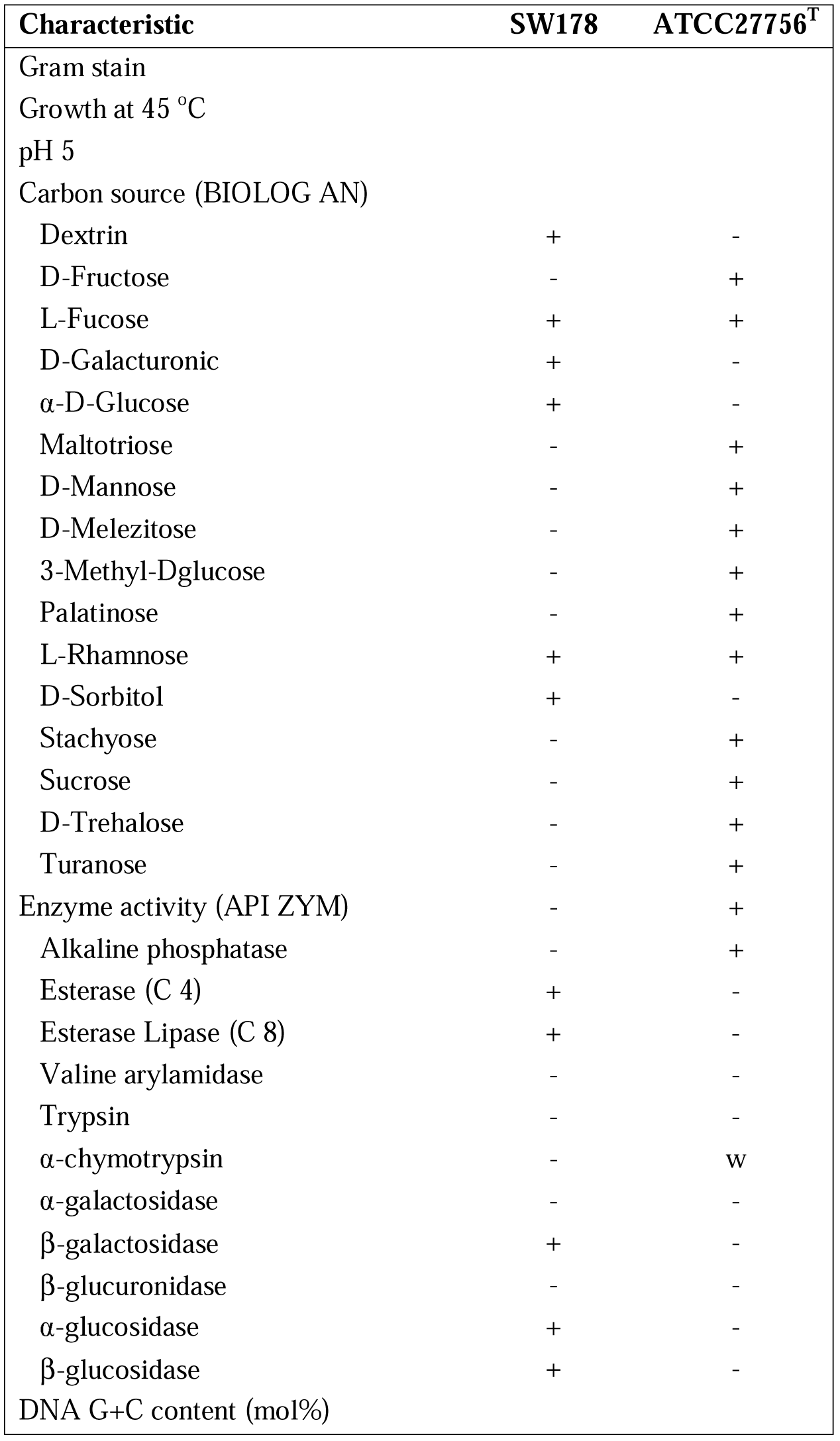
Characteristics of *R. catenae* sp. nov. (= CCOS 1886 ^T^, =DSM 109242^T^) and the closest related strain. Strains SW178 and *Ruminococcus torques* ATCC 27756^T^ were anaerobically cultured in BHIM at 37 °C for 2 days. +, Positive; -, negative; w, weakly positive; ND, not detected.

| Characteristic | SW178 | ATCC27756<sup>T</sup> |
| --- | --- | --- |
| Gram stain |  |  |
| Growth at 45 °C |  |  |
| pH 5 |  |  |
| Carbon source (BIOLOG AN) |  |  |
| Dextrin | + | - |
| D-Fructose | - | + |
| L-Fucose | + | + |
| D-Galacturonic | + | - |
| α-D-Glucose | + | - |
| Maltotriose | - | + |
| D-Mannose | - | + |
| D-Melezitose | - | + |
| 3-Methyl-Dglucose | - | + |
| Palatinose | - | + |
| L-Rhamnose | + | + |
| D-Sorbitol | + | - |
| Stachyose | - | + |
| Sucrose | - | + |
| D-Trehalose | - | + |
| Turanose | - | + |
| Enzyme activity (API ZYM) | - | + |
| Alkaline phosphatase | - | + |
| Esterase (C 4) | + | - |
| Esterase Lipase (C 8) | + | - |
| Valine arylamidase | - | - |
| Trypsin | - | - |
| α-chymotrypsin | - | w |
| α-galactosidase | - | - |
| β-galactosidase | + | - |
| β-glucuronidase | - | - |
| α-glucosidase | + | - |
| β-glucosidase | + | - |
| DNA G+C content (mol%) |  |  |

**Table 2.** Cellular fatty acid contents (%) of strain *R. catenae* sp. nov. (= CCOS 1886 ^T^, =DSM 109242^T^) and its closest related strain. Strains SW178 and *Ruminococcus torques* ATCC 27756^T^ were anaerobically cultured for 2 days in BHIM. Fatty acids comprising less than 1% of the total fatty acids in both strains is not shown.

| Fatty acid composition | SW178 | ATCC27756<sup>T</sup> |
| --- | --- | --- |
| **Straight chain** |  |  |
| C<sub>12:0</sub> | 1.31 | 2.9 |
| C<sub>14:0</sub> | 28.71 | 7.98 |
| C<sub>16:0</sub> | 19.55 | 23.58 |
| C<sub>16:0</sub> ALDE | 3.35 | 2.46 |
| C<sub>18:0</sub> | 1.15 | 9.96 |
| **Demethylacetal (DMA)** |  |  |
| C<sub>12:0</sub> DMA |  |  |
| C<sub>14:0</sub> DMA | 7.82 | 0.78 |
| C<sub>16:0</sub> DMA | 12.55 | 9.66 |
| C<sub>18:0</sub> DMA | 1.73 | 3.42 |
| C<sub>16:1</sub>ω7c DMA | 2.41 | - |
| C<sub>18:1</sub>ω7c DMA | 7.9 | - |
| **Unsaturated** |  |  |
| C<sub>13:1</sub> at 12-13 | 4.26 | 0.38 |
| C<sub>16:1</sub>ω7c | 1.42 | 10.26 |
| C<sub>16:1</sub>ω9c | - | 6.15 |
| C<sub>18:1</sub>ω7c | 1.95 | 3.57 |
| C<sub>18:1</sub>ω9c | 0.78 | 16.71 |
| Summed Feature 1 | 4.26 | 0.38 |

Based on the genotypic and phenotypic features of strains SW178 and *Ruminococcus torques* ATCC 27756 ^T^, we propose that strain SW178 should be described as a new member of the genus Ruminococcus. We assign strain SW178 as the type strain of the new species, for which the name *Ruminococcus catenae* is proposed.

## Description of *Ruminococcus catenae* SW178 sp. nov

*Ruminococcus catenae* SW178 sp. nov. (L. Pl. n. *catenae*, referring to cells growing in chains). The strain is strictly anaerobic, Gram-strain-positive and non-motile. The average size of each cell is 0.5-1.0 µm, oval-shaped, and growing in chains. Colonies are visible on BHI-M agar after 2 days and are approximately 0.1–0.3 cm in diameter, white, raised and circular with entire edge. The microorganism exhibits optimal growth in BHI-M medium at 45° C and pH 7.5. The strain utilizes dextrin, L-fucose, D-galacturonic, α-D-Glucose, L-rhamnose and D-sorbitol. The primary cellular fatty acids are C_14□:□0_, C_16□:□0_ and C_16□:□0_ DMA. The genome of this strain is 3.18 Mbp with 46.9 mol% of G+C content. This strain was isolated from the cecum of feral chicken. The type strain is SW178 (= CCOS 1886 ^T^, =DSM 109242^T^)

## Protologue

The 16S rRNA gene and genome sequence were deposited in GenBank under accession number MN133850 and PRJNA554364, respectively. The digitized protologue of the strain SW178 is TA01018.

## Acknowledgments

This work was supported in part by the USDA National Institute of Food and Agriculture, Hatch projects SD00H532-14 and SD00R540-15, and a grant from the South Dakota Governor’s Office of Economic Development awarded to JS. SW received support from the Science Achievement Scholarship of Thailand.

